# CoaTran: Coalescent tree simulation along a transmission network

**DOI:** 10.1101/2020.11.10.377499

**Authors:** Niema Moshiri

## Abstract

**Motivation:** The ability to simulate coalescent viral phylogenies constrained by a given transmission network can enable the benchmarking of computational tools used in molecular epidemiology as well as the ability to gain insights into unobservable aspects of the virology of a novel pathogen. However, such simulation experiments require generating a large number of technical simulation replicates, and existing tools for coalescent simulations along a transmission network are too slow to conduct such experiments at the scale of the global population.

**Results:** CoaTran is a massively scalable tool that simulates a coalescent viral phylogeny constrained by a user-provided transmission network. CoaTran is written in highly-optimized C++ code and can generate global population scale phylogenetic coalescent simulations in seconds to minutes.

**Availability:** CoaTran is freely available at https://github.com/niemasd/CoaTran as an open-source software project.

**Supplementary information:** Supplementary data are available online.

## 1 Introduction

Modern methods of molecular epidemiology typically attempt to summarize or reconstruct properties of a viral outbreak from a multiple sequence alignment (Pond *et al*., 2018) or phylogeny (Balaban *et al*., 2019; Ragonnet-Cronin *et al*., 2013; Prosperi *et al*., 2011) inferred from genome sequences of viral samples collected from patients. However, because the true transmission history of a real outbreak is typically not known or is error-prone, the accuracies and errors of such methods can be assessed in simulation (Moshiri *et al*., 2018). Further, the ability to simulate realistic social contact networks, transmission networks, viral phylogenies, and sequences modeled after real-world outbreaks can provide insights into the unobservable aspects of the virology of a novel pathogen (Worobey *et al*., 2020).

Given known sample times from a simply-modeled population, the coalescent (Kingman, 1982) provides an elegant and efficient method for simulating a phylogeny: begin with each sample as a leaf lineage, and iteratively “coalesce” two lineages until only a single lineage remains. Coalescent simulation becomes slightly more complicated when accounting for continuous time rather than discrete time as well as when allowing effective population size to vary over time (e.g. exponential or logistic effective population growth).

The simulation of a viral phylogeny can be framed in a coalescent framework: given a transmission network (either estimated from real-world data or simulated under a compartmental epidemiological model) and given collection times for each viral sample, simulate a coalescent phylogeny with each viral sample as a leaf, but constrain the phylogeny such that each transmission dictates a bottleneck event. Inferences made from such simulations are generally strengthened as more simulation replicates are generated, yet existing methods of coalescent simulation along a transmission network are prohibitively slow for global population scale simulation studies.

Here, we introduce CoaTran, an open-source tool that simulates a coalescent viral phylogeny constrained by a user-provided transmission network. Notably, CoaTran is written in highly-optimized C++ code to minimize memory allocations/deallocations and to maximize memory locality. As a result, CoaTran is orders of magnitude faster than the only known existing method, and it enables global population scale phylogenetic coalescent simulations in seconds to minutes.

## 2 Related work

To our knowledge, the only existing tool that allows users to simulate a coalescent phylogeny constrained by a given transmission network is the VirusTreeSimulator component of the PANGEA.HIV.sim package (Ratmann *et al*., 2016), which was the motivation for the development of CoaTran. With respect to within-host effective viral population size, VirusTreeSimulator allows users to simulate phylogenies under constant effective population size as well as exponential and logistic effective population growth from the time of infection. In its current form, CoaTran only supports coalescence with constant effective population size, which is reasonable for many real-world contexts (Worobey *et al*., 2020), and we plan to incorporate exponential and logistic effective population growth in a future update. However, CoaTran is consistently ~100x faster than VirusTreeSimulator (Fig. 1), making it the appropriate tool for global population scale studies. Note that both CoaTran and VirusTreeSimulator run single-threaded.

**Fig. 1.**
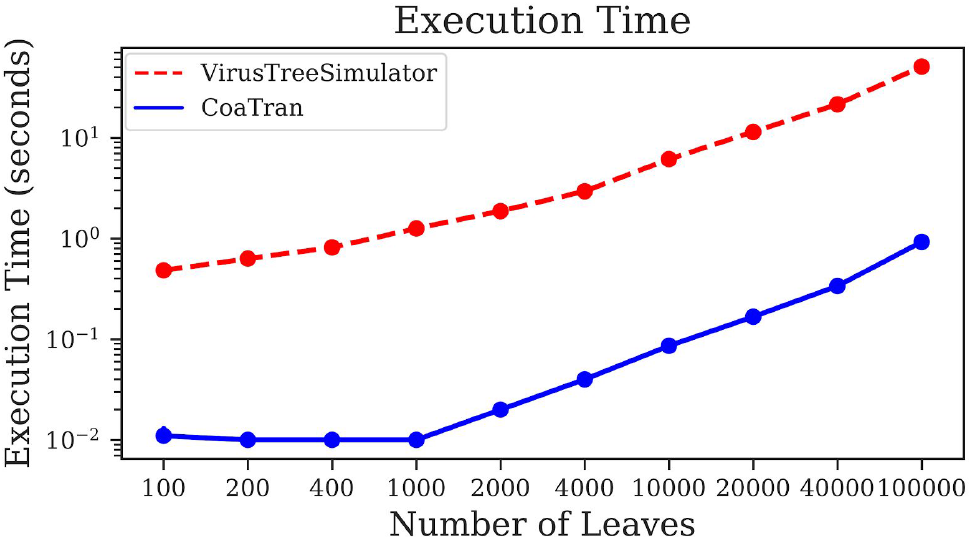
Execution time. Execution time of VirusTreeSimulator and CoaTran for phylogenies of varying sizes. Each point is the average of 10 technical replicates. The lack of visible error bars is due to the small variance in runtime across replicates. All runs were executed sequentially on a single core of a 1.8 GHz Intel i7-8565U CPU.

## 3 Results and discussion

CoaTran is written in C++, and the code base aims to minimize memory allocations/deallocations and to maximize memory locality for speed. CoaTran has no external library dependencies and can be easily compiled in any C++ environment with `make`, making it user-friendly to install as well as portable across platforms. CoaTran takes the following as input: (1) a transmission network, (2) the patient sample times, and (3) the effective population size of the coalescent process, and it outputs a simulated coalescent tree in the Newick format. If there are multiple transmission chains in the transmission network, a Newick tree will be output corresponding to each transmission chain.

The execution times of VirusTreeSimulator and CoaTran were compared using simulated epidemic datasets. First, a contact network was simulated under the Barabási–Albert (BA) model (Barabási & Albert, 1999) using NetworkX (Hagberg *et al*., 2008). The contact network had 100,000 individuals and had an expected degree of 4 contacts per individual. Next, an epidemic was simulated along the contact network under the Susceptible–Infected (SI) compartmental epidemiological model using GEMF (Sahneh *et al*., 2017) until the desired number of infected individuals (i.e., number of leaves in the phylogeny) was reached. A per-infected-neighbor infection rate of 0.1 was arbitrarily chosen to yield reasonably large amounts of time between consecutive transmission events to avoid the bounds of floating point numerical precision. A single sample time was randomly selected for each infected individual uniformly along the window of time between the individual’s initial infection and twice the overall outbreak simulation time; this choice was arbitrarily made to yield reasonably long terminal branches in the phylogeny. A constrained coalescent phylogeny was then simulated using both VirusTreeSimulator and CoaTran. This process was repeated for each technical simulation replicate, and there were a total of 10 simulation replicates for each desired number of leaves.

As can be seen in Figure 1, CoaTran is consistently orders of magnitude faster than VirusTreeSimulator: even on outbreak simulations with 100,000 infected individuals, for which VirusTreeSimulator takes approximately one minute to complete per technical replicate, CoaTran is able to complete in less than one second. This difference may seem small in a single execution, but it is important to note that simulation experiments that utilize tools like VirusTreeSimulator and CoaTran can require thousands or even millions of simulation replicates to sufficiently support the findings, thus compounding the speedup significantly.

As mentioned earlier, CoaTran currently only supports coalescence with constant within-host effective viral population size, but we aim to implement exponential and logistic population growth (and potentially other models of population size over time) in future updates. Further, although CoaTran currently only runs single-threaded, the algorithm it employs can natively parallelize across multiple transmission chains in a single epidemic simulation as well as across multiple parallel paths in a single transmission chain, and we aim to implement multithreading in a future update.

## Acknowledgements

We would like to thank Jonathan Pekar and Joel Wertheim for fruitful discussions, and we would like to thank Matthew Hall for his excellent write-up of the mathematical derivations behind continuous-time coalescent simulations.

